# Immunovisualization of spatial changes in leaves and root tissue associated with drought stress in wheat (*Triticum aestivum* L.)

**DOI:** 10.1101/2025.10.06.680837

**Authors:** Agata Leszczuk, Nataliia Kutyrieva-Nowak, Tomasz Skrzypek

## Abstract

**Background and Aims:** Plants have evolved complex cell-type-specific processes to adapt to a dynamic environment, exhibiting distinct signals in response to emerging drought stress. We propose an advanced qualitative and quantitative analysis approach, demonstrating tissue specificity in drought adaptation, which in turn may provide novel biological insights.

**Methods:** We performed immunofluorescence labeling of specific cellular components *in situ*, and the acquired data were analyzed in terms of changes in quantitative and spatial fluorescence intensity.

**Results:** The qualitative analysis revealed differences in terms of individual components and individual days of the experiment. The quantitative analysis of leaf anatomy showed that the most pronounced changes were observed in the level of proteoglycans (JIM13, JIM15) and polysaccharides (LM5, LM16, LM20). The leaves of plants growing in drought were characterized by destroyed fragments, in which increased secretion of extensins, AGPs, galactans, hemicelluloses, and RG-I was noted. In turn, the qualitative analyses of the microscopy images of roots, along with fluorescence intensity analyses, revealed a significantly higher content of AGP and arabinoxylan in the exodermis in plants grown under drought stress.

**Conclusion:** Our research has revealed that the changes at the tissue level are targeted and highly specific. The obtained results also emphasize the importance of *in planta* analyses, which indicate that findings from only single *ex planta* studies may distort the entire image of changes occurring in the plant as a result of stress.

**HIGHLIGHT STATEMENT:** One of the strategies employed by plants to mitigate the effects of water loss is the mechanical protection of organs through targeted and highly specific modifications in their cellular architecture.

## INTRODUCTION

Climate change is reducing water availability, negatively affecting plant survival and biodiversity conservation worldwide. In turn, biodiversity loss may be critical to the capacity of agriculture to provide sufficient crops for the ever-growing population and, consequently, threaten food security. Thus, early and effective detection of water stress in plants is essential to avoid losses in the ecosystem, emphasizing the need to develop methodologies and strategies to cope with drought stress (Munné-Bosch and Villadangos 2023; Wang et al. 2024). Plants have evolved complex cell-type-specific processes to adapt to a dynamic environment, exhibiting distinct signals in response to emerging drought stress (Gupta et al. 2020; Cantó-Pastor et al. 2024).

Drought stress causes a range of structural changes in plants at the cellular, tissue, and organ levels (Bhanbhro et al. 2024). Changes in the leaf include e.g. guard cell shrinking, stomatal aperture reduction, a decrease in turgor pressure in bulliform cells, epidermal shrinkage, and an increase in cutin and wax deposition. The changes in the root include an increased root-to-shoot ratio, increased lignification, and strengthening of the xylem structure (Verelst et al. 2013; Ouyang et al. 2020; Ganie and Ahammed 2021). All these changes are the result of changes at the molecular level, which in turn disrupt the proper differentiation of the tissue or become an adaptation that allows survival in unfavorable water-deficient conditions. Also, the reduction in metabolic costs aims to improve water transport efficiency, thereby enhancing drought tolerance (Fonta et al. 2022). One of the primary effects is a reduction in water content within cells, which is associated with loss of turgor pressure and decreased cell expansion. Also known are the processes of plasmolysis and cell membrane breakdown as well as signal transduction, which links water deficiency to developmental reprogramming. To counteract osmotic stress, plants accumulate proline and soluble sugars, as well as lignin and suberin, which help stabilize cell structures and maintain osmotic balance (Ezquer et al. 2020; Ganie and Ahammed 2021; Bhanbhro et al. 2024). Based on all the observed quantitative changes in the assembly of plant cells under drought stress, several potential mechanisms have been proposed that govern adaptive responses allowing the plant to reduce their metabolic demand and protect vital cellular functions in drought conditions (Moore et al. 2008; Tenhaken 2015). However, the information on the key structural alterations occurring within plant cells under drought stress remains limited; in particular, there are no detailed data on the differentiation between root-specific and leaf-specific adaptive responses.

Our previous reports indicate that the biochemical composition of the wheat plant cell is susceptible to stress, probably constituting a mechanism of the plant response to adaptation to unfavorable environmental conditions (Leszczuk and Kutyrieva-Nowak 2025). We concluded that water loss can be linked to changes in the content of pectins and their ability to interlink with ions, which influences the elemental economy due to modifications in the cellular assembly. Previous analyses allow us to hypothesize that wall stiffening is connected with drought tolerance and turgor maintenance (Leszczuk and Kutyrieva-Nowak 2025). Although there is a substantial amount of data on the changes occurring in wheat plants under drought stress, from advanced omics studies (Bhanbhro et al. 2024; Guo et al. 2025) to comprehensive physiological analyses (Ru et al. 2025), the spatial patterns of tissue alterations remain inconsistent and lack uniform characteristics. Therefore, the current study builds upon our earlier molecular studies exploring plant variability in response to drought stress. Here, we performed imaging of immunolocalization mapping of specific cellular components *in situ*, and the acquired data were analyzed in terms of changes in quantitative and spatial fluorescence intensity. Using a standard method, we propose an advanced qualitative and quantitative analysis approach, demonstrating tissue specificity in drought adaptation, which in turn may provide novel biological insights. In this comparative study, all data obtained from the material subjected to drought stress are compared with data from the control material (well-watered plants). The first step was to analyze the general fluorescence intensity. Then, we compared the changes within two experimental terms, i.e. after 5 and 20 days of drought. The analyses showed that the quantitative changes were not always synchronous with the microscopic images viewed. Hence, in the next step, tissue specificity in the plant response to stress was demonstrated, because the fluorescence intensity was not uniform throughout the entire tissue. In this way, we were able to show that plants respond to stress in a very selective and complex way. The obtained results also emphasize the importance of *in planta* analyses, which indicate that findings from only single *ex planta* studies may distort the entire image of changes occurring in the plant as a result of stress.

## MATERIAL & METHODS

### Pot experiment

The experiment was carried out in pots (10 cm x 10 cm) filled with soil taken from a wheat field (51°25′53.5” N, 19°37′05.3” E). The research material was 5-day-old winter wheat seedlings (*Triticum aestivum*) cv. ‘Arkadia’, (named: start–day 0), which were then subjected to drought stress. Then, after 5 days and 20 days of the experiment, leaves and roots were collected, and the preparative part for microscopic analyses began. A control experiment, in which the seedlings were regularly watered, was carried out in parallel. All seedlings were grown at 23°C under a 16-hour light/8-hour dark cycle. The experiment was conducted on 50 seedlings per sample variant, from which chemically fixed cubes of research material were obtained. Immunocytochemical reactions with all antibodies were performed on consecutive sections of the same block of research material.

### Immunofluorescence technique

Excised seedling fragments were fixed in 2% (w/v) paraformaldehyde (PFA, Sigma) and 2.5% (v/v) glutaraldehyde (GA, Sigma) in phosphate-buffered saline (PBS, Sigma) under vacuum for 2 h. Next, the material was rinsed in PBS (three times, 15 min each) and the dehydration process (in a graded series of ethanol solutions from 30%, 50%, 70%, 90%, to 99.8% for 20 min) was carried out at room temperature. Ethanol was substituted with 3:1, 1:1, and 1:3 mixtures of EtOH and LR White resin (Sigma) for 2 hours and with pure LR White resin overnight. The polymerization in gelatin capsules was performed at 55 °C for 48 h. Semithin (1 µm) sections were obtained using an ultramicrotome (PowerTome XL, RMC Boeckeler, USA) equipped with a glass knife and were mounted on poly-L-lysine-coated glass slides. Slides prepared for immunochemistry reactions were circled with a liquid blocker PAP Pen (Sigma).

Sections adhering to the poly-L-lysine slides were washed twice in PBS and then treated with 1% bovine serum albumin (BSA) in PBS for 30 minutes to prevent nonspecific binding of antibodies. Then, the sections were incubated with the primary antibody present at a 1:50 dilution in 0.1% BSA at 4 °C for 24 h and washed four times with PBS. The experiment was conducted using monoclonal antibodies obtained from Kerafast collections (USA). Table 1 describes the primary monoclonal antibodies used in immunolabeling procedures. Importantly, for visualization of the examined epitopes, goat anti-rat IgM conjugated with AlexaFluor 488 was used as a secondary antibody (Thermo Fisher Scientific, USA). The secondary antibody was diluted 1:200 in 1% BSA at 4 °C and left in the dark. After 24 h, the incubated sections were washed in PBS and deionized water, and finally enclosed in Dako Fluorescent Mounting Medium (Sigma).

**Table 1.**
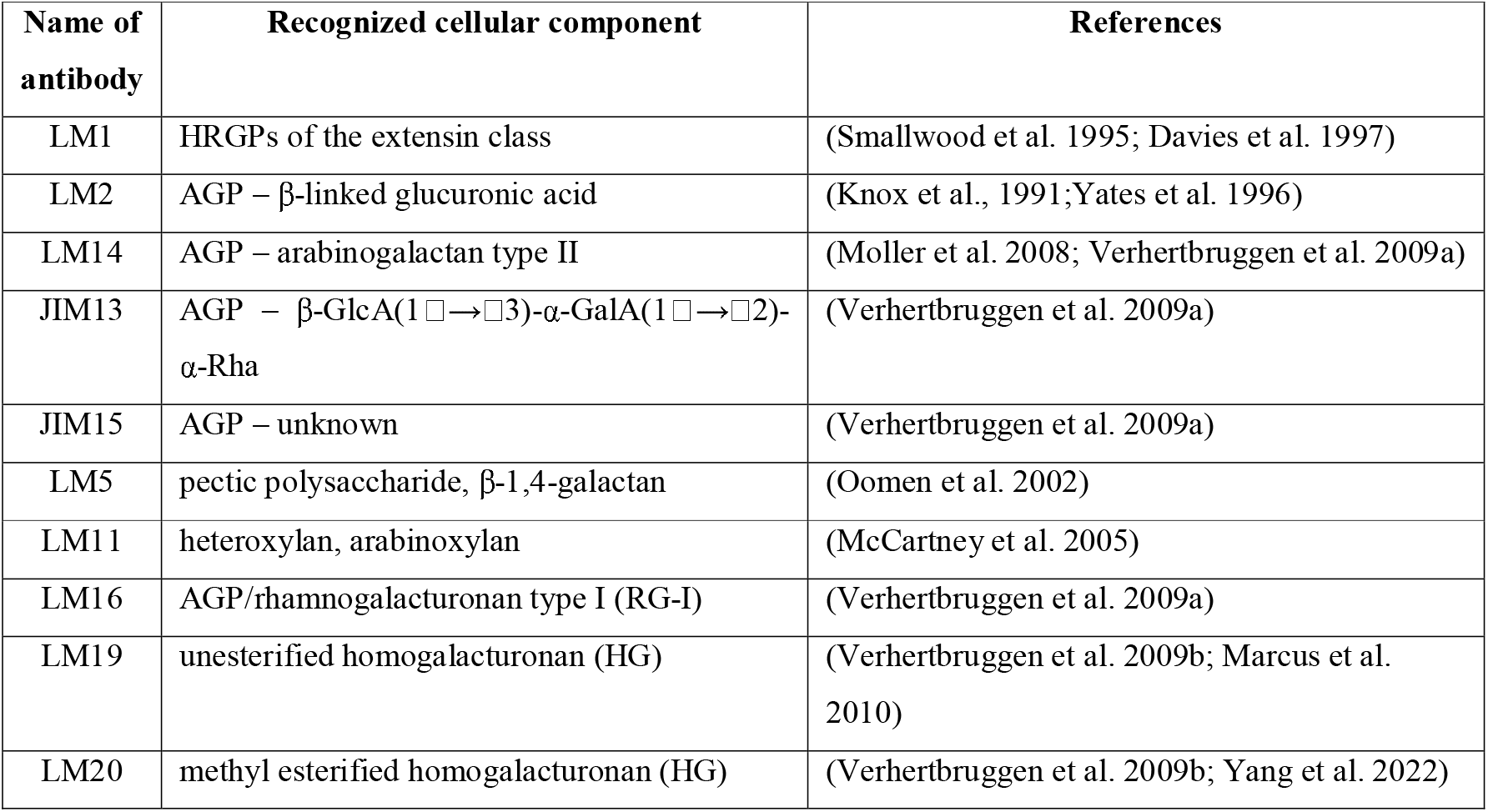
Description of antibodies used in the current studies.

### Imaging and fluorescence intensity analyses

The observations and imaging were carried out using a Confocal Laser Scanning Microscopy (ZEISS Axio Observer Z1, LSM 700, Jena, Germany) coupled with ZEN Standard software. All parameters (i.e., laser intensity, gain) were kept constant for all samples. The excitation wavelength for AlexaFluor488 was 490 nm, and the emission was collected at 525 nm. Control reactions were carried out by omitting the primary antibody. Immunofluorescence labeling was carried out on at least several serial sections from each sample and each kind of antibody. During imaging, 5 images were taken from a single sample. Representative image sets were selected and edited using the CorelDRAW X6 graphics program. Average fluorescence pixel intensity was measured with the Zeiss ZEN software and ImageJ 1.51 software. From the obtained data, the arithmetic mean of the fluorescence intensity from the green channel (fluorescence indicating the presence of the antibody) was calculated. For the analysis of a particular tissue in the same section, individual measurements were performed using the fluorescence intensity value at a specific signal dot. The data were statistically analyzed using Statistica v.13 tools (TIBCO Software Inc., USA). For comparisons of the mean values, an analysis of variance (one-way ANOVA), followed by a Tukey post hoc honestly significant difference test, was used. For all analyses, the significance level was estimated at p < 0.05.

## RESULTS

### Structural changes in leaves during drought

The first stage of the study involved the qualitative assessment of changes in the structural arrangement of individual cell components. The structural elements analyzed can be divided into the following categories: proteins (LM1), proteoglycans (LM2, JIM13, JIM15, LM14), polysaccharides (LM5, LM16, LM19, LM20), and hemicelluloses (LM11). Already the 5-day drought caused morphological and anatomical changes in the leaf tissue, with a noticeably distorted tissue arrangement. The volume of the cells decreased as a result of water deprivation for 5 days, leading to general shrinkage and turgor loss in the leaf tissue. After 20 days of drought, the leaf was destroyed with numerous modifications. None of these changes were visible in the leaves of the control plants (Figure 1).

**Figure 1.**
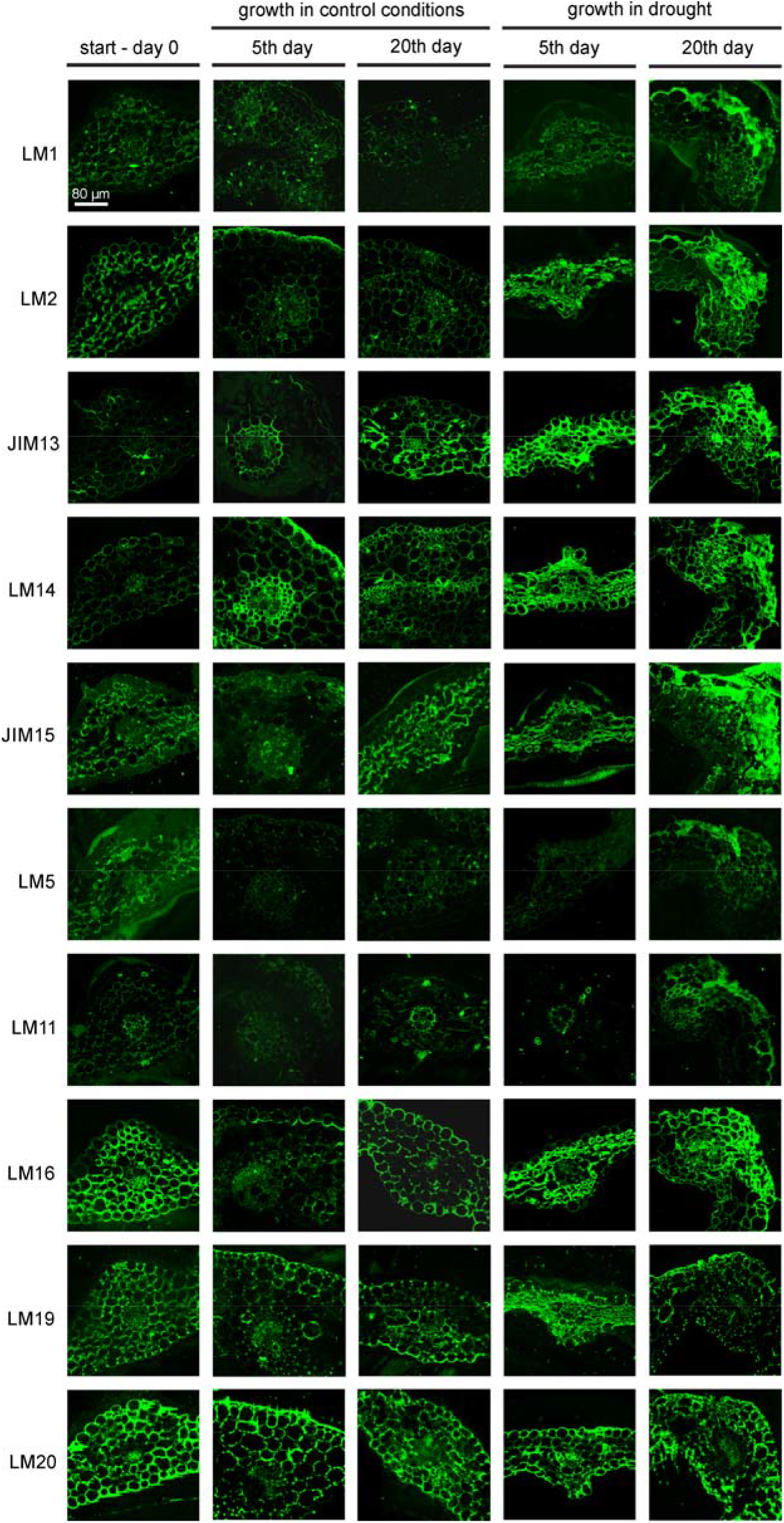
Representative images showing spatial changes in the distribution of cellular components in wheat leaves cultivated in drought conditions for 5 and 20 days. The first column represents images of seedling leaves at the start of the experiment (day 0), followed by images of seedling leaves from the watered cultivation. Immunolocalization of extensin (LM1), AGPs (LM2, JIM13, LM14, JIM15), galactans (LM5), xylans (LM11), AGPs/RG-I (LM16), and HGs (LM19, LM20). CLSM imaging. Green fluorescence indicates a positive reaction and the presence of a specific epitope. The level of the reaction intensity is directly correlated with the varying levels of compound accumulation. Bars for all photographs: 80 µm.

The identification of the cell components showed that not all of them responded to the drought conditions in the same way. Moreover, differences were also noted between the leaves of the 5-day-old seedlings (day 0) and the control plants and plants grown in the drought conditions. The qualitative analysis revealed differences in terms of individual components and individual days of the experiment. While extensins (LM1) did not have a characteristic pattern, AGPs always constituted a boundary in the cell wall-cell membrane continuum. In turn, the analyzed polysaccharides (LM19, LM20) were mainly present in the cell wall, filling most of its space, and in intercellular junctions. A significantly lower amount of galactans (LM5) and hemicelluloses (LM11) was also observed, in comparison to the other labeled components (Figure 1).

Based on the obtained images, a quantitative analysis of the structural changes in the leaf tissue was conducted. The quantitative analysis of fluorescence intensity showed that the 5-day drought did not cause statistically significant spatial changes in the distribution of all components (Figure 2). Interestingly, after 20 days of growth in the drought conditions, structural reorganization occurred, but not in all cases. The overall quantitative analysis of the leaf anatomy images showed the most pronounced changes in the level of proteoglycans (JIM13, JIM15) and polysaccharides (LM5, LM16, LM20). Another observation is the increased dynamics of changes in proteoglycans, here exemplified by AGP. Its presence in the drought conditions changed significantly, compared to its presence in the leaf tissue of the watered plants. The amount of AGP recognized by JIM13 was increased in the leaves of plants grown in the drought conditions, while the level of AGP recognized by JIM15 decreased almost 4-fold. The analyses of changes in polysaccharides showed that the fluorescence intensity of galactans (LM5), RG-I (LM16) and methyl esterified HGs (LM20) increased, relative to the control samples (Figure 2).

**Figure 2.**
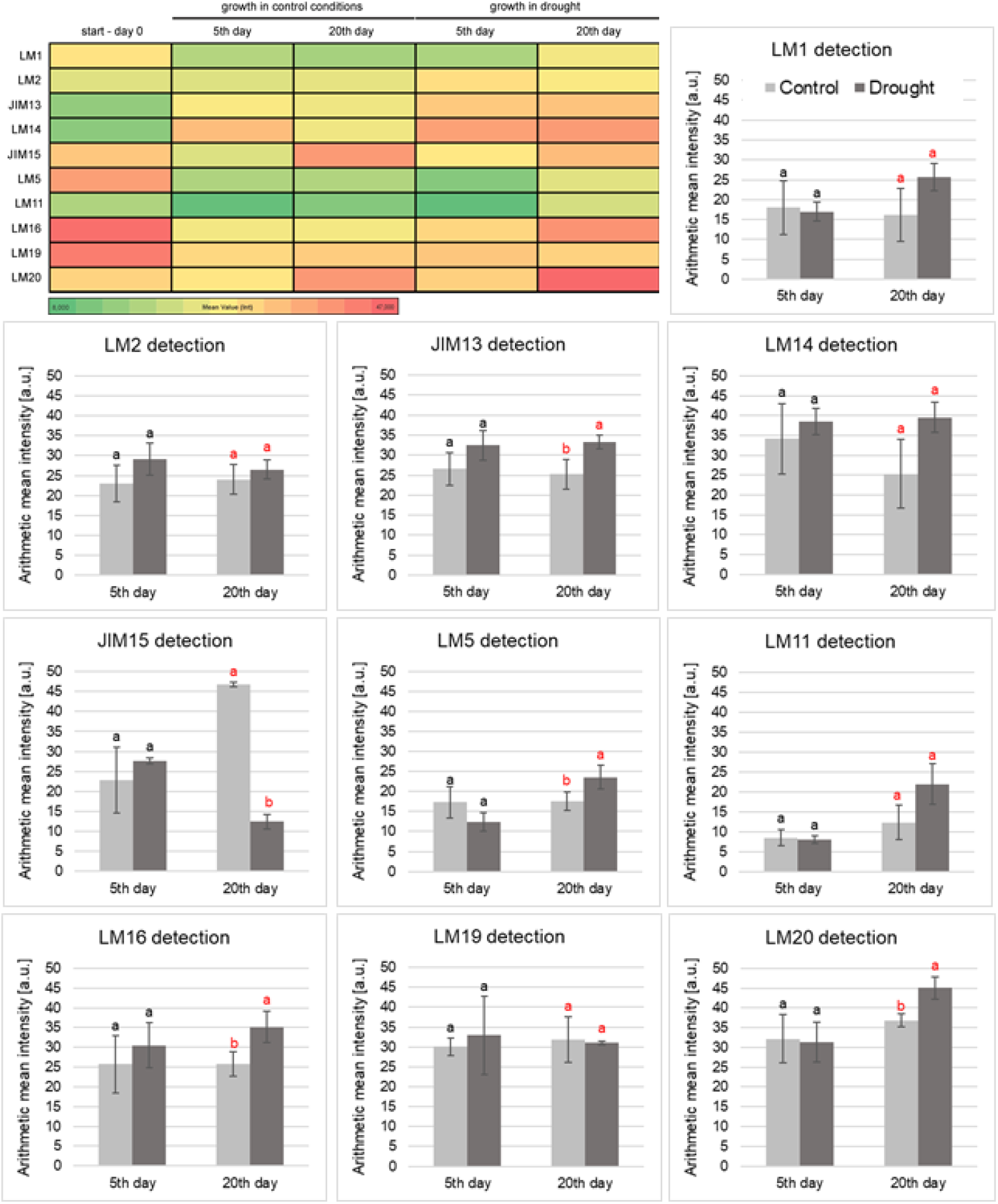
Analysis of fluorescence intensity in sections of wheat leaves grown in drought conditions for 5 and 20 days. Comparative analysis with control samples - watered wheat. Immunolocalization of extensin (LM1), AGPs (LM2, JIM13, LM14, JIM15), galactans (LM5), xylans (LM11), RG-I (LM16), and HGs (LM19, LM20). The overall visualization is a heatmap that represents the intensity of fluorescence across the samples. The color intensity corresponds to the magnitude of the data, with red colors indicating higher values and green colors representing lower values. Statistical significance in the graphs was determined using a Tukey post-hoc test (different letters indicate a significant difference at p < 0.05).

The detailed analysis of the spatial distribution of labeled components often showed visible accumulation of larger amounts of components in the changed parts. This concerned the external leaf compartments in plants growing for 5 days in the drought conditions. The accumulation of AGPs identified by antibodies JIM13, LM14, and LM2 was mainly observed. In turn, the leaves of plants growing under drought for 20 days were characterized by destroyed fragments, in which increased secretion of extensins (LM1), AGPs (LM2, JIM15), galactans (LM5), hemicelluloses (LM11), and RG-I (LM16) was also noted. The aforementioned components filled nearly the entire damaged and deformed cells, as shown in Figure 3, where leaf fragments with accumulation are marked by white arrows (for 5 days) and a red dotted line (for 20 days). Additionally, the intensity value profile plots from the degraded tissue often show a fluorescence signal that is multiple times higher, specifically in the damaged areas of the leaf tissue (Figure 3).

**Figure 3.**
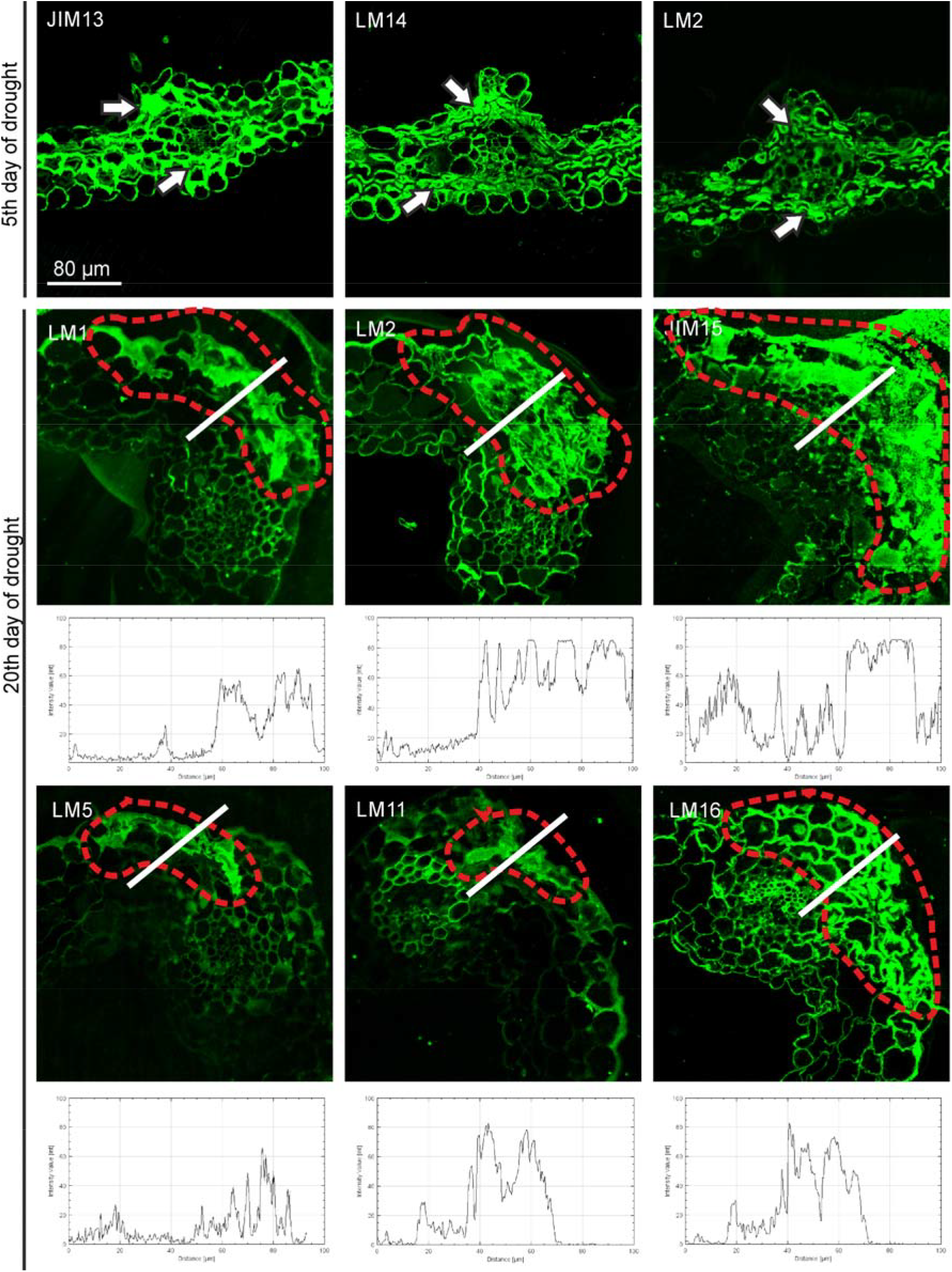
Representative images with spatial-structural changes in leaves in response to drought stress. Selected images showing the most prominent changes in the structure of leaf tissue and plots of fluorescence intensity values in the selected areas (white line). Marked spots with the highest intensity in the modified parts of the leaf after 5 days of drought (white arrows) and after 20 days of drought (red dotted lines). Immunolocalization of components with the most significant spatial changes (JIM13, LM14, LM2, LM1, JIM15, LM5, LM11, LM16). CLSM imaging. Bars for all photographs: 80 µm.

### Structural changes in roots during drought

The anatomical root analysis showed that the root of the 5-day-old wheat seedling (day 0) was not yet fully differentiated. In turn, the roots of the plants after the next 5 days of the experiment had typical root tissues, i.e., the epidermis, cortex, and vascular bundles (Figure 4). Similarly, the labeling of the cellular components showed differences between the root of the 5-day-old seedling and the older roots. In the root of the 5-day-old seedling, there was no typical arrangement of cellular components, while clear spatiotemporal differences in their arrangement were visible in the roots after 5 and 20 days of the experiment. The components identified are representative of different root tissues. The highest diversity was visible in the case of the AGP distribution, because its various epitopes were visible in different root structures. β-linked glucuronic acid (LM2) and arabinogalactan type II (LM14) were visible in the vascular bundle, but β-GlcA(1□→□3)-α-GalA(1u→u2)-α- Rha (JIM13) was present in all root tissues. In turn, JIM15 was almost invisible. Hemicelluloses exhibited a very characteristic and selective distribution, which was noted only in the pericycle and endodermis. In turn, homogalacturonans were almost not visible, which indicates their low content in the root tissue (Figure 4).

**Figure 4.**
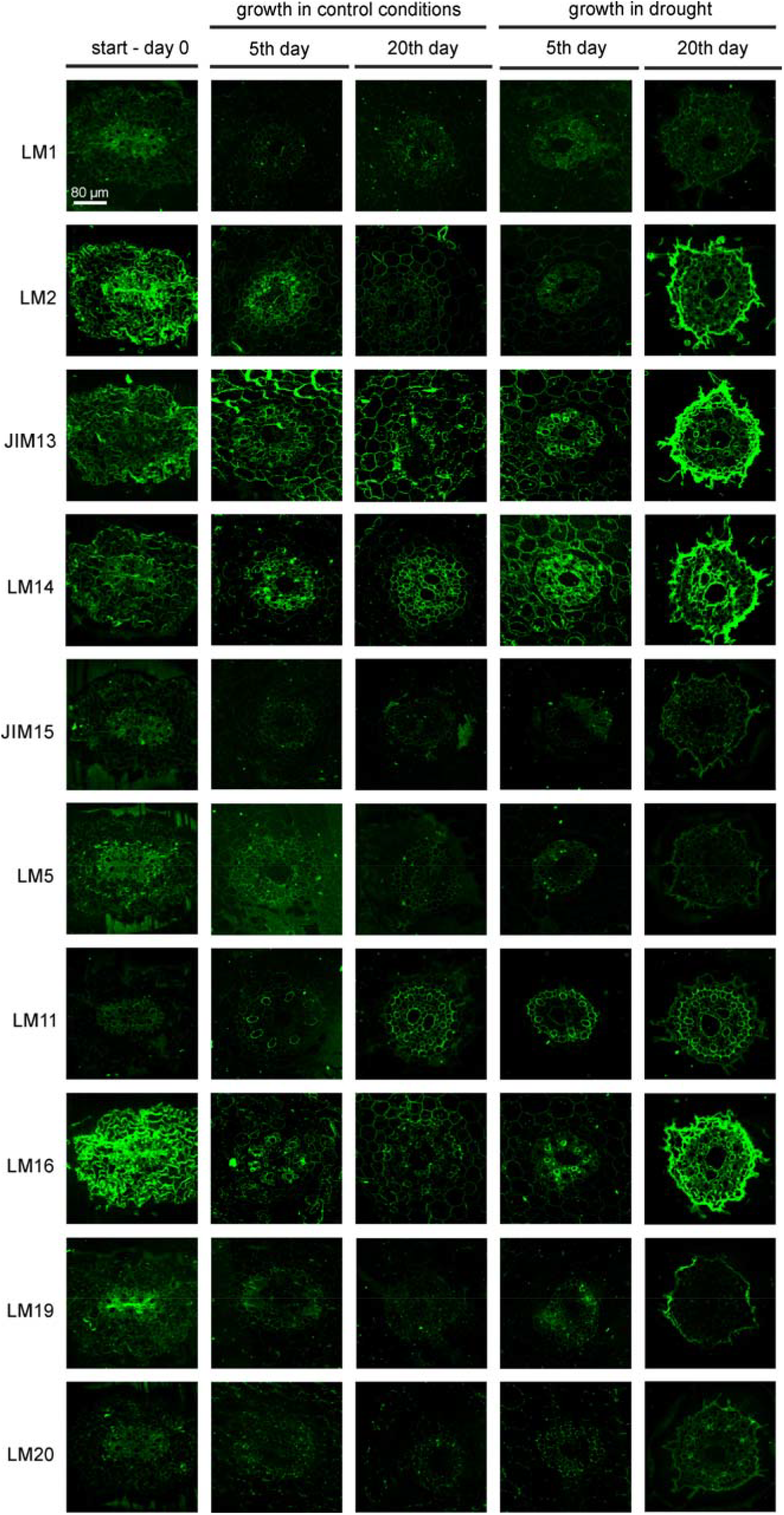
Spatial changes in the distribution of cellular components in wheat roots cultivated in drought conditions for 5 and 20 days. The first column represents images of seedling roots at the start of the experiment (day 0), followed by images of seedling roots from the watered cultivation. Immunolocalization of extensin (LM1), AGPs (LM2, JIM13, LM14, JIM15), galactans (LM5), xylans (LM11), RG-I (LM16), and HGs (LM19, LM20). CLSM imaging. Green fluorescence indicates a positive reaction and the presence of a specific epitope. The level of the reaction intensity is directly correlated with the varying levels of compound accumulation. Bars for all photographs: 80 µm.

The qualitative analysis showed that the drought conditions affected the root structure and potentially altered the distribution of individual cell components, but only the quantitative fluorescence measurements facilitated precise determination of spatial changes. The differences in the cellular arrangement after 5 and 20 days of growth in the drought conditions were indicated by the qualitative analysis of the microscopic images (Figure 4). They were confirmed by the fluorescence intensity measurements in roots cultivated in the water deficit conditions for 5 days, which showed a significant decrease in the content of AGP (LM2, JIM13), galactans (LM5), and RG-I (LM16). In turn, the comparative analysis of the root after 20 days of drought showed a significant increase in the fluorescence intensity of AGP (LM2, JIM13, LM14) and RG-I (LM16). The aforementioned increase was often twofold larger in the roots of plants growing in drought. In the case of detection of JIM15 and LM5, a significant decrease in fluorescence intensity was noted. Interestingly, no differences were noted for HG (LM19, LM20) and hemicelluloses (LM11), and their fluorescence intensity levels indicated that they were present in trace amounts (Figure 5).

**Figure 5.**
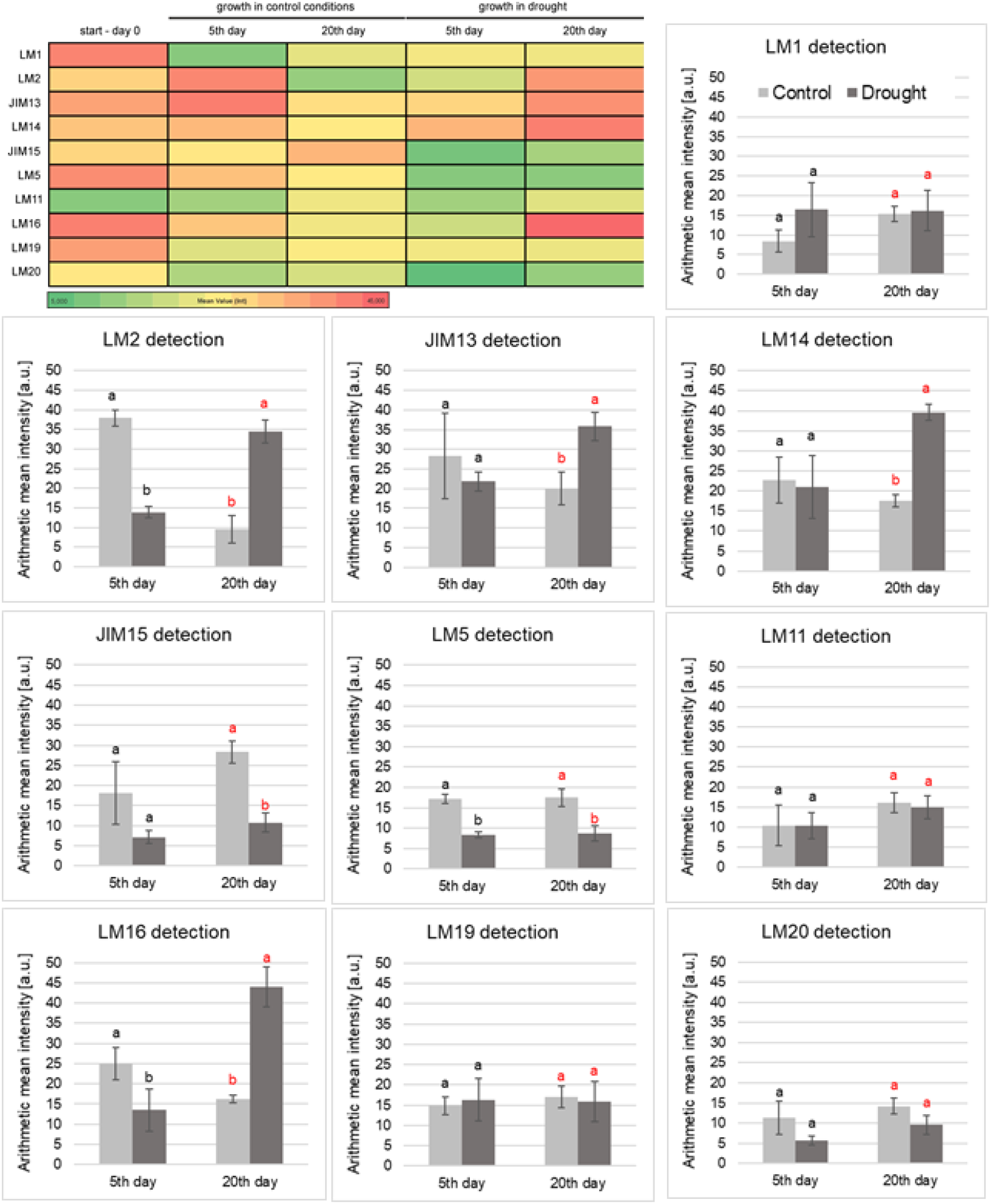
Analysis of fluorescence intensity in sections of wheat roots grown in drought conditions for 5 and 20 days. Comparative analysis with control samples - watered wheat. Immunolocalization of extensin (LM1), AGPs (LM2, JIM13, LM14, JIM15), galactans (LM5), xylans (LM11), RG-I (LM16), and HGs (LM19, LM20). The visualization is a heatmap that represents the intensity of fluorescence across the samples. The color intensity corresponds to the magnitude of the data, with red colors indicating higher values and green colors representing lower values. Statistical significance was determined using a Tukey post-hoc test (different letters indicate a significant difference at p < 0.05).

The fluorescence intensity analysis showed that, in the case of AGP and RG-I, the measured signal was significantly higher, but only in the outer part of the root – the exodermis of the wheat plants after 20 days of drought. The fluorescence intensity measurement in this tissue was 5 times higher compared to the other root tissues. Additionally, the intensity value profile plots from the outer part of the roots confirmed the significantly higher fluorescence signal (Figure 6). It can be estimated that the outer part of the root, characterized by the presence of secretion, forms an approximately 20-µm thick layer. Moreover, no such difference was noted in the roots of plants in the 5-day experiment. Interestingly, no increased accumulation was observed in the case of extensins, galactans, hemicelluloses, and HGs.

**Figure 6.**
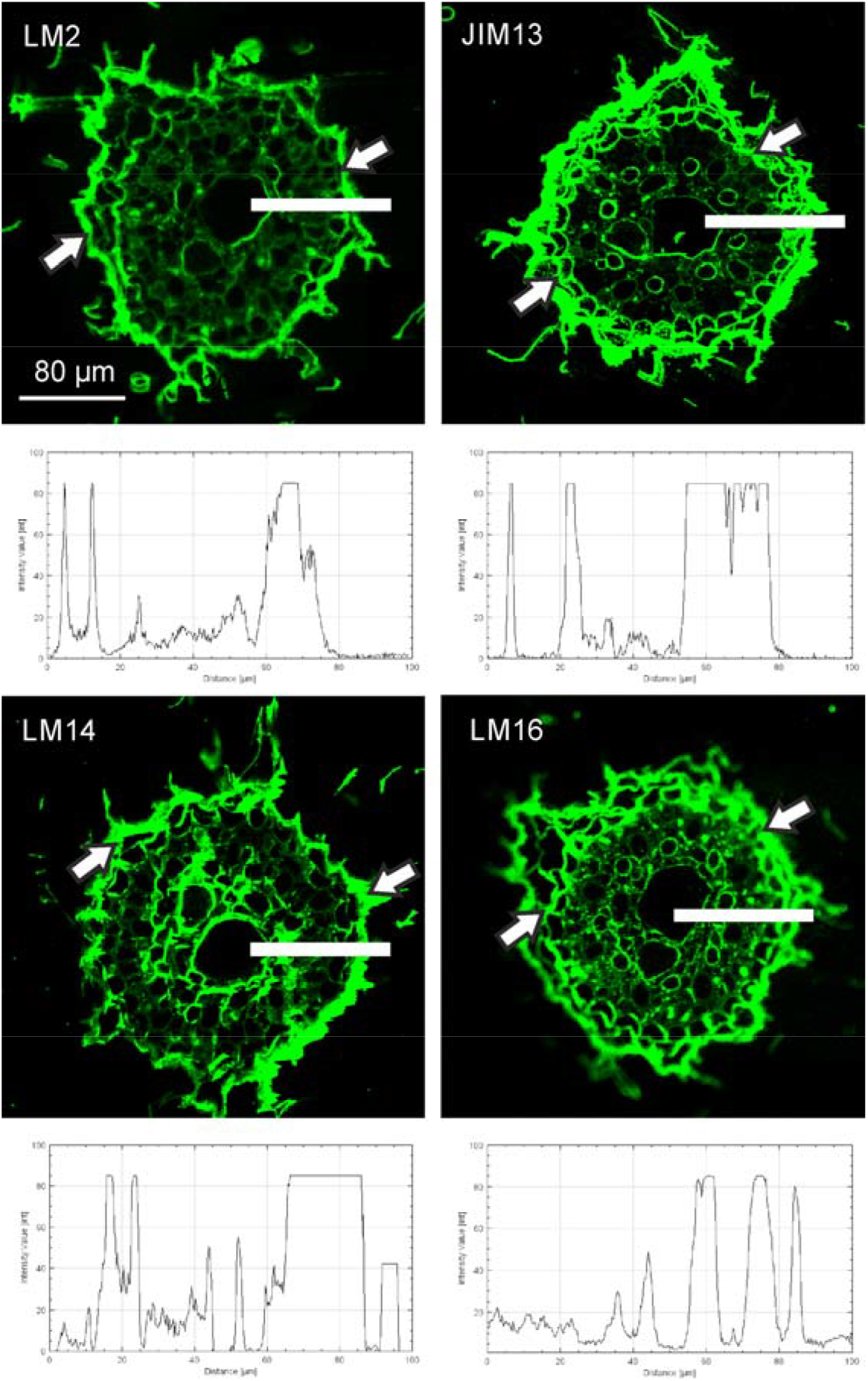
Representative images with spatial-structural changes in roots in response to drought stress. Selected images showing the most prominent changes in the structure of root tissue and plots of fluorescence intensity values in the selected areas (white line). Marked spots with the highest intensity in the modified external parts of the roots after 20 days of drought (white arrows). Immunolocalization of components with the most significant spatial changes (LM2, JIM13, LM14, LM16). CLSM imaging. Bars for all photographs: 80 µm.

## DISCUSSION

The research presented in the current paper encompassed two key aspects: a scientific one and a technical one. An exceptionally relevant topic regarding the adaptive changes in plants in response to water deficit caused by the increasing occurrence of drought conditions was addressed. The research problem was approached using tools that are typically not the primary choice for such analyses, specifically the immunolocalization method for visualizing *in planta* changes. The next step involved consideration of quantitative changes based on microscopy images. Microscopic studies reveal spatial differentiation and the presence of specific secretion, which cannot be investigated through analyses based on tissue homogenization and extraction of individual components. Our analyses indicate that the chosen approach is indeed the most appropriate and necessary for a comprehensive understanding of plant adaptations to growth under abiotic stress.

In the experiment, a wheat seedling was exposed to drought stress. The stress duration encompassed two distinct periods: 5 days of drought, which is a commonly observed phenomenon in recent years, and 20 days of drought, a critical period that occurs sporadically and often results in catastrophic effects on crops. The drought periods were chosen based on our previous biochemical analyses, in which we demonstrated the onset of the first changes/responses (5 days) and the moment leading to complete tissue necrosis (20 days) (Leszczuk and Kutyrieva-Nowak 2025). Here, we analyzed the structural changes both in the roots, as the first organs exposed to water deficit (Shimazaki et al. 2005), and in the leaves, as they are crucial for such processes as photosynthesis and other related functions. In our study, we obtained clear evidence that structural modifications in root cells occur by the 5th day of drought, whereas in leaves, these changes become visible much later, specifically after 20 days. Moreover, both the qualitative and quantitative analyses indicated changes within the root and leaf tissues; however, they represented entirely different structural modifications. It was noted that the water deficit resulted in morphological modifications and numerous cell deformities. In these altered parts of the organs, increased secretion of certain components was observed, i.e. polysaccharides in the case of the leaves. In the case of the roots, elevated secretion of AGPs was observed, primarily in the outer regions of the root, where the main secretion filling the cells of the exodermis was formed.

An interesting aspect of the conducted analyses is the identification of high tissue specificity related to the plant response to growth in drought stress conditions. Quantitative changes in the cell assembly are often not fully compatible with *in situ* qualitative studies due to the specific secretion within individual cellular compartments. In general, the secretion of substances outside the root structures is a well-known adaptive mechanism (Ouyang et al. 2020). Ouyang et al. (2020) focused on the adaptive responses of roots to water deficit stress, specifically the selective accumulation of suberin and lignin in the exodermis - the outermost layer of the organ. Using rice and wheat as examples, the authors explain that suberin seals the cells, thereby limiting transpiration, while lignin provides mechanical support, preserving the structural integrity of the roots (Ouyang et al. 2020). Similarly, analyses of tomato plants grown in drought stress conditions revealed increased secretion in the exodermis, which was explained as an apoplastic diffusion barrier regulating the flow of water (Cantó-Pastor et al. 2024). External root layer secretions may also serve as a protective barrier, as supported by the role of pectic side chains during the water stress response in a drought-tolerant wheat cultivar, as shown by Leucci (2008). Our qualitative analyses of the microscopy images, along with the fluorescence intensity analyses, revealed a significantly higher content of AGP and arabinoxylan in the roots of plants grown under drought stress. It is important to highlight that the exodermis in the root showed the highest fluorescence intensity, indicating marked accumulation of these compounds. In the drought conditions, the mucilage production increased as part of the plant response. We assume that AGPs in the mucilage help maintain hydration around the root zone by forming a sticky gel that retains moisture. This finding aligns with previous reports on the role of arabinogalactans of cell wall modifications in drought adaptation (Moore et al. 2008; Nguema-Ona et al. 2013; Tenhaken 2015; Ezquer et al. 2020). It is assumed that high arabinan levels contribute to maintaining the fluidity of the pectin network in the cell wall during desiccation (Jones et al. 2003; Moore et al. 2008; Tenhaken 2015). Morre et al. (2008) suggest that arabinose polymers act as ‘pectic plasticizers’ and may play a role in maintaining the flexibility of cell walls during dehydration. Structural changes involve modifying the balance between wall extensibility and turgor pressure, directly affecting the non-covalent intra- and interpolymeric bonds within the wall due to water removal.

In the case of the leaves, the spatial changes at the tissue level appeared later than in the roots. The comparative analysis suggests that a different rearrangement of the cell structural components occurs in the leaves, compared to the roots. In the drought conditions, the fluorescence intensity measurements in the leaf tissues indicated the most pronounced signal associated with the accumulation of pectic polysaccharides, specifically, β-1,4-galactan, RG-I, and methyl esterified HG. The secretion of pectic compounds in leaf tissue modifications may indicate their role in protection. Our findings are consistent with the work by Leucci et al. (2008), in which the mol% of side chains of RG I and II significantly increased in response to water stress in “drought-tolerant” wheat seedlings cv. Capeiti (Leucci et al. 2008). Another confirmation is the studies conducted by Jones et al. (2003), who proposed that arabinans play a crucial role in determining the physical and functional properties of guard cell walls. Transpiration depends on the opening and closing of stomata, which in turn is influenced by the degree of expansion of the leaf guard cells (Lin et al. 2022). Alterations in arabinan content can influence the regulation of stomatal dynamics, particularly by affecting stomatal opening and closure. Furthermore, the authors suggested a mechanism in which arabinans contribute to maintaining cell wall flexibility by preventing HG polymers from forming tight associations, thereby contributing to greater plasticity in the cell wall structure (Jones et al. 2003).

Additionally, it is important to focus on the anatomy and morphology of leaves in plants grown in drought conditions. These leaves exhibit extensively damaged tissues, which may indicate the occurrence of the Programmed Cell Death (PCD) process. As is well-established, PCD is a regulated mechanism that enables plants to cope with drought stress by eliminating damaged or non-functional cells, thereby potentially protecting the entire plant. During drought stress, PCD in leaves may play a crucial role in preventing excessive water loss by selectively removing compromised cells. In the research conducted by Hameed et al. (2013), two wheat genotypes with differing drought tolerance levels were examined to assess PCD induction in leaves. The study revealed that the occurrence of PCD correlated with enhanced activities of peroxidase, superoxide dismutase, and catalase as well as elevated ascorbate levels in drought stress conditions (Hameed et al. 2013). Furthermore, in our studies, heightened secretion was detected in the damaged regions of the leaf, occupying the degenerated cells. The fluorescence intensity analyses demonstrated differences in the signal in the plants exposed to the 20-day drought stress, specifically reflecting an increase in fluorescence intensity in reactions with antibodies labeling AGP, galactans, and RG-I. In the study on cell wall remodeling under stress, it is suggested that the increase in side chains of pectic polysaccharides may arise from the formation of hydrated gels by pectins, which serve to mitigate cellular damage (Tenhaken 2015). In turn, the accumulated presence of AGPs enables optimal mechanical integrity by creating a “buffer zone” that prevents direct interaction between cell membranes and the cell wall matrix (Mareri et al. 2019; Ezquer et al. 2020). Interestingly, studies carried out to identify and analyze the expression of AGPs in rice roots have revealed genes that are differentially expressed in drought-tolerant and drought-sensitive genotypes (Rabello et al. 2008).

Taken together, these results form the basis for further scientific investigations. In our work, we have demonstrated that one of the strategies that plants employ to cope with the effects of water loss is the mechanical protection of organs by modifying their cell/cell wall architecture. Our research has revealed that the changes at the tissue level are targeted and highly specific. In turn, these efforts pave the way for devising a genetic engineering approach for developing drought-tolerant wheat plants. Importantly, plant varieties with an increased level of arabinans/AGP, which serves as a protective secretion in the roots, could be developed. Furthermore, focus could be placed on varieties with higher content of leaf polysaccharides, which potentially play an important role in general mechanisms of maintaining the mechanical strength and flexibility of the cell wall under drought. Moreover, the obtained data on the native adaptation of plants to drought form the basis for studies on changes in soil properties. The addition of soil conditioners with a protective role against desiccation remains an unresolved issue. The present data on structural changes could serve as an indicator of the action of soil modifiers, which would not only alter soil properties but also positively impact plant growth.

## STATEMENTS & DECLARATIONS

### Author Contributions

A.L. Conceptualization, Investigation, Visualization, Writing – original draft; N.K. Visualization, Writing - review and editing; T.S. Investigation, Writing - review and editing

### Data Availability

The datasets generated during the current study are available from the corresponding author on reasonable request.

